# Two birds with one stone: a novel potential antibiotic blocking IsdB-mediated heme extraction by *Staphylococcus aureus* with serendipitous hemoglobin left-shifting activity

**DOI:** 10.1101/2025.05.29.656759

**Authors:** Sarah Hijazi, Francesco Marchesani, Marialaura Marchetti, Valeria Buoli Comani, Paul Brear, Barbara Campanini, Luca Ronda, Serena Faggiano, Eleonora Gianquinto, Somayeh Asghar Pour Hassan Kiyadeh, Barbara Rolando, Francesca Spyrakis, Carlotta Compari, Loretta Lazzarato, Omar De Bei, Emanuela Frangipani, Stefano Bettati

**Author notes:** Contributed equally to the work (Sarah Hijazi, Francesco Marchesani, Marialaura Marchetti).

## Abstract

Infections caused by *Staphylococcus aureus* are closely linked to its ability to secure essential nutrients, including iron, which is extracted from the heme of human hemoglobin (Hb) through the iron-regulated surface determinant (Isd) system. The compound 4-[[2-[[5- (1H-indol-3-yl)-1,3,4-oxadiazol-2-yl]sulfanyl]acetyl]amino]benzoate (C35) was recently identified as a new potential antimicrobial agent for its ability to bind Hb and hamper its interaction with the staphylococcal hemophore IsdB *in vitro*. Here, we show that C35 inhibits *S. aureus* growth by specifically targeting the hemophore-driven iron acquisition system. Our findings confirm both the potential of C35 as a first-in-class protein-protein interaction inhibitor with antimicrobial activity, and the effectiveness of targeting hemophores as a strategy to inhibit *S. aureus* growth. To gain information for drug discovery purposes, the X-ray structure of Hb in the presence of the compound was solved. Unexpectedly, we discovered that, rather than the predicted binding pose, the molecule binds to tetrameric Hb in a cleft between the alpha subunits, stabilizing an R2 relaxed Hb conformation. This triggered further investigation of the effect of C35 on Hb functional properties, which showed a pronounced left-shift activity on oxygen binding curve (i.e., it strongly increases the Hb oxygen affinity). These results highlight C35 as a promising dual-acting compound with both antimicrobial activity and the ability to modulate Hb function through non-covalent stabilization of a high-affinity state.

**Author Summary:** *Staphylococcus aureus* is a dangerous bacterium that can cause severe infections in humans. To grow and survive it needs iron, which it steals from our red blood cells by taking it from hemoglobin, the protein that carries oxygen in the blood. In this study, we focused on a small molecule, called C35, that blocks the interaction between hemoglobin and a key bacterial protein involved in heme acquisition. We found that C35 strongly inhibits the growth of *S. aureus* when hemoglobin is the only available source of iron, showing a potential new method to starve the pathogen and consequently fight the infection. Surprisingly, we also found that C35 increases the affinity of hemoglobin for oxygen. This dual action makes C35 a unique molecule for future therapeutic development, with potential applications both as a new antimicrobial agent and in the treatment of diseases related to hemoglobin function.

## Introduction

*Staphylococcus aureus* is a human commensal and opportunistic pathogen capable of causing severe infections. Its ability to develop resistance to antibiotics highlights the urgent need for new, effective antimicrobial agents [1]. *S. aureus* depends on iron for survival; in the human body, this metal is primarily associated with heme cofactor in hemoglobin (Hb). With the aim of gaining access to the hemic iron, during infection *S. aureus* induces red blood cell (RBC) hemolysis exploiting hemolysins, thus allowing access to free Hb [2]. Iron gain is mediated by the iron-regulated surface determinant (Isd) system, a nine-protein gear in charge of intercepting Hb, extracting heme and internalizing it [3]. During the first step, the cell wall-anchored hemophore IsdB interacts with circulating Hb [4] and extracts the heme [5], which is transferred to other proteins of the Isd system in order to complete the iron acquisition process [2]. IsdB is a multifaceted protein that plays a crucial role in *S. aureus* virulence by facilitating iron acquisition, promoting adherence to host tissues, and modulating the host immune response [6–9]. Due to its significant role in *S. aureus* pathogenesis, IsdB has been explored as a potential vaccine candidate. However, clinical trials have proven unsuccessful, underscoring the need for innovative antibacterial strategies to combat *S. aureus* infections [10]. One promising approach is the discovery of chemical entities able to counteract the IsdB:Hb complex formation [11]. With this purpose, in a recent work we conducted a virtual screening campaign targeting the IsdB:Hb complex, and *in vitro* binding tests led to the identification of 4-[[2-[[5-(1H-indol-3-yl)-1,3,4-oxadiazol-2-yl]sulfanyl]acetyl]amino]benzoate (C35) as the most potent molecule, with a K_D_ for Hb of 0.57 ± 0.06 µM [12]. In this work, we performed microbiological tests that proved the C35 effectiveness in inhibiting *S. aureus* growth in the presence of Hb as the sole iron source. We also solved the crystallographic structure of Hb in complex with C35 to provide the basis for chemical optimization of the compound. The structural analysis revealed a binding pose different from the one initially hypothesized [12] but similar to that of a known class of molecules which allosterically increase the affinity of Hb for oxygen by stabilizing its relaxed, high-affinity state. These molecules are known as left shifters, as they shift the Hb oxygen binding curve leftwards, i.e. to lower partial oxygen pressures. Under a pharmacological perspective, the biological effect of left shifters is primarily exploited in the treatment of sickle cell disease (SCD), where the stabilized state delays the polymerization of sickle Hb (HbS) [13]. This finding prompted us to investigate the effect of C35 on increasing oxygen affinity of Hb, a property that could confer an additional functional role to the molecule.

## Results

### C35 inhibits S. aureus growth in the presence of Hb as sole iron source

In a recent study, we have identified a compound named C35 that binds Hb with high affinity (K_D_ of 0.57 ± 0.06 µM) and interferes with the formation of the IsdB:Hb complex [12]. These results suggested that C35 could act as a promising antimicrobial agent capable of disrupting iron acquisition in *S. aureus*, thereby leading to bacterial iron starvation and consequent growth inhibition. To investigate this hypothesis, a mutant lacking the IsdB component of the Isd system was generated in *S. aureus* Newman (i.e., *S. aureus* Δ*isdB*). Since the expression of the Isd system is regulated by the Ferric Uptake Regulator (Fur), which enables its expression under iron-poor conditions, the chemically-defined medium NRPMI devoid of iron was used [14]. To identify the best condition for C35-inhibitory testing, we first investigated the role of Hb in supporting *S. aureus* growth in our experimental setting, using both the WT and the Δ*isdB* strains. Bacteria were pre-cultured in RPMI supplemented with 500 µM EDDHA, to iron-starve bacterial cells. The following day, cultures were sub-inoculated in NRPMI in the presence of 500 µM EDDHA, supplemented or not with either 80 nM Hb or 500 µM FeCl_3_, and monitored overtime for 96 h (Fig 1A). Results showed that both strains failed to grow in the absence of an iron source, indicating that EDDHA, when added at 500 µM, effectively prevents the acquisition of iron traces present in NRPMI, which would likely occur through the production of siderophores, usually expressed under iron-depleted conditions [15]. Interestingly, supplementation with 80 nM Hb promoted the growth only of the WT strain but not of the Δ*isdB* (Fig 1A), in line with previous findings [6,14]. Notably, this Hb-mediated growth promotion was observed after 96 h post-incubation, suggesting that *S. aureus* requires time to sense the presence of Hb in the medium, and subsequently expresses the IsdB hemophore. In contrast, both strains were able to readily utilize FeCl_3_, showing comparable growth, thus confirming the specificity of IsdB for Hb utilization (Fig 1A). Altogether, these data highlight the phenotypic characterization of the Δ*isdB* mutant, thus providing a foundation for subsequent Hb-inhibitory testing.

**Fig 1.**
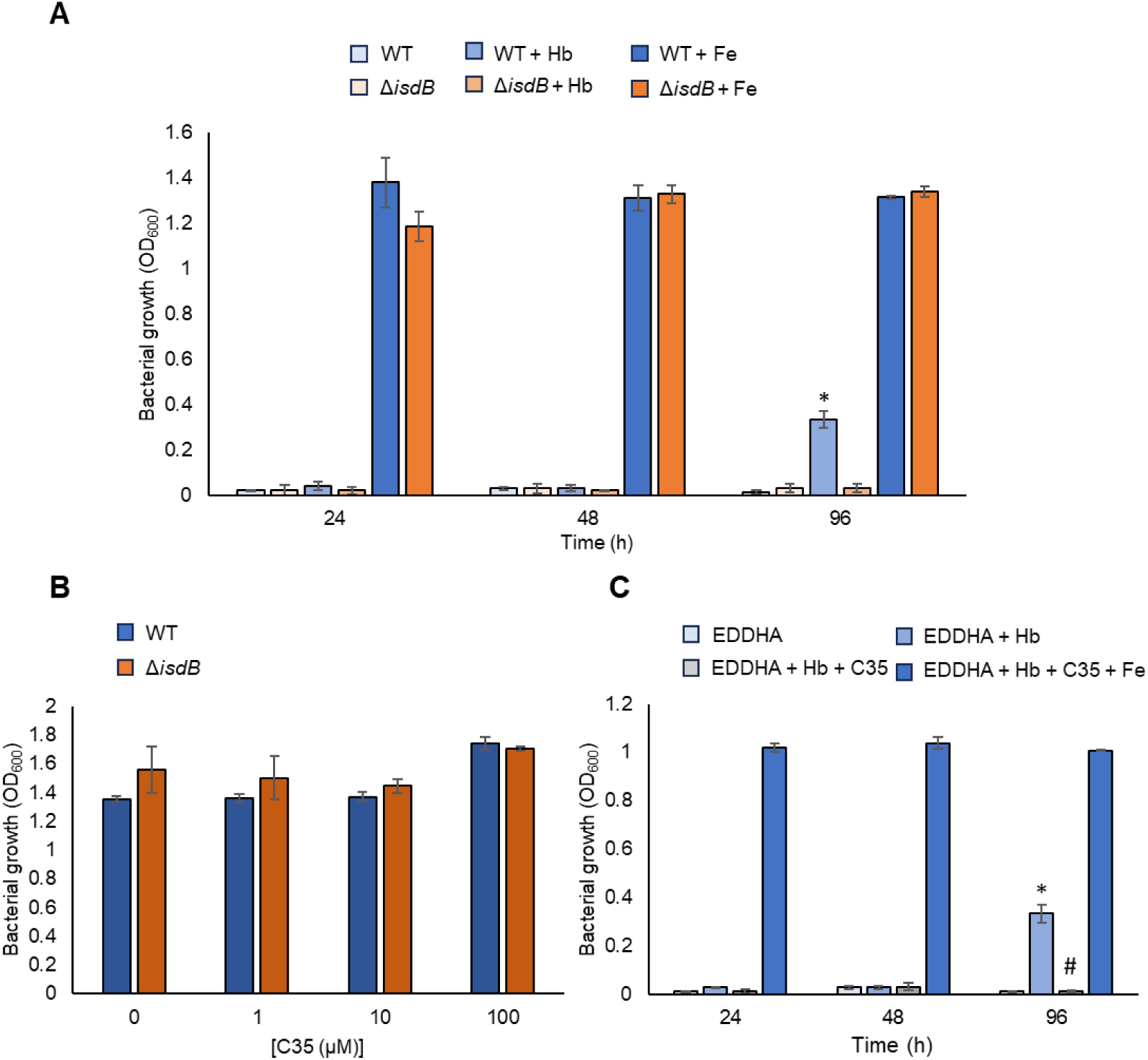
C35 inhibits *S. aureus* growth in the presence of Hb as the sole iron source. *S. aureus* WT and its isogenic Δ*isdB* mutant were pre-cultured in RPMI supplemented with 500 µM EDDHA, to restrict iron availability, for 16-20 h at 37 °C with good aeration (shaking 180 rpm). Next day, cultures were washed and resuspended in NRPMI supplemented or not with the indicated compounds. (A) Effect of 80 nM Hb (Hb) and 500 µM FeCl3 (Fe) on *S. aureus* WT and Δ*isdB* growth, in RPMI + 500 µM EDDHA; (B) Toxicity of C35 on *S. aureus* WT and Δ*isdB*, grown for 24 h in NRPMI without EDDHA, to allow the growth of both strains; (C) Effect of 100 µM C35 on *S. aureus* WT grown in NRPMI + 500 µM EDDHA (EDDHA) supplemented with 80 nM Hb alone or in combination with 500 µM FeCl3. As negative control, the growth of the WT strain in NRPMI + 500 µM EDDHA was also included. Each value is the average of three different cultures ± standard deviation. The symbols indicate statistically significant differences as determined by a Student’s t test (P < 0.05) in relation to the WT strain grown in the presence of EDDHA with no Hb (*) or the WT strain grown in the presence of EDDHA with Hb (#).

C35 toxicity was investigated by monitoring the growth of WT and Δ*isdB* in NRPMI without EDDHA, to allow bacterial growth, in the presence of different C35 concentrations (i.e., 1, 10 and 100 µM), and compared to the one of the positive control, devoid of the compound (Fig 1B). No growth defect was observed for both strains at all C35 concentrations tested, compared to the unamended controls (Fig 1B). Following the toxicity assessment of C35 towards *S. aureus*, the maximum concentration used (100 µM) was then selected to investigate the ability of this compound to inhibit Hb-mediated growth promotion in the WT strain. To this aim, *S. aureus* WT cells were cultivated in NRPMI supplemented with 500 µM EDDHA and/or Hb, in the presence or absence of C35. Interestingly, C35 completely inhibited the WT growth in the presence of 80 nM Hb as the sole iron source after 96 h post-incubation, thus indicating that it is able to block IsdB-mediated Hb uptake in *S. aureus* (Fig 1C), while supplementation with 500 µM FeCl₃ as an alternative iron source, completely abolished C35-mediated growth inhibition (Fig 1C). Additional evidence supporting the proposed mechanism of action confirmed that C35 exerts its inhibitory effect by binding to Hb, making it selective for the hemophore-mediated iron acquisition system. Indeed, isothermal titration calorimetry (ITC) measurements excluded any measurable interaction between C35 and IsdB at the tested concentrations (S1 Fig).

### Three-dimensional structure of Hb-C35 complex

Given the ability of C35 of inhibiting *S. aureus* growth and aiming to further explore this molecule as a potential lead compound, we decided to investigate the binding of C35 to Hb at atomic resolution. The X-ray crystallographic structure of tetrameric Hb in complex with C35 was obtained by co-crystallization (PDB ID: 9HBA and Fig 2A). The final resolution was 1.5 Å and the crystals form showed a symmetry belonging to the space group P 32 2 1.

**Fig 2.**
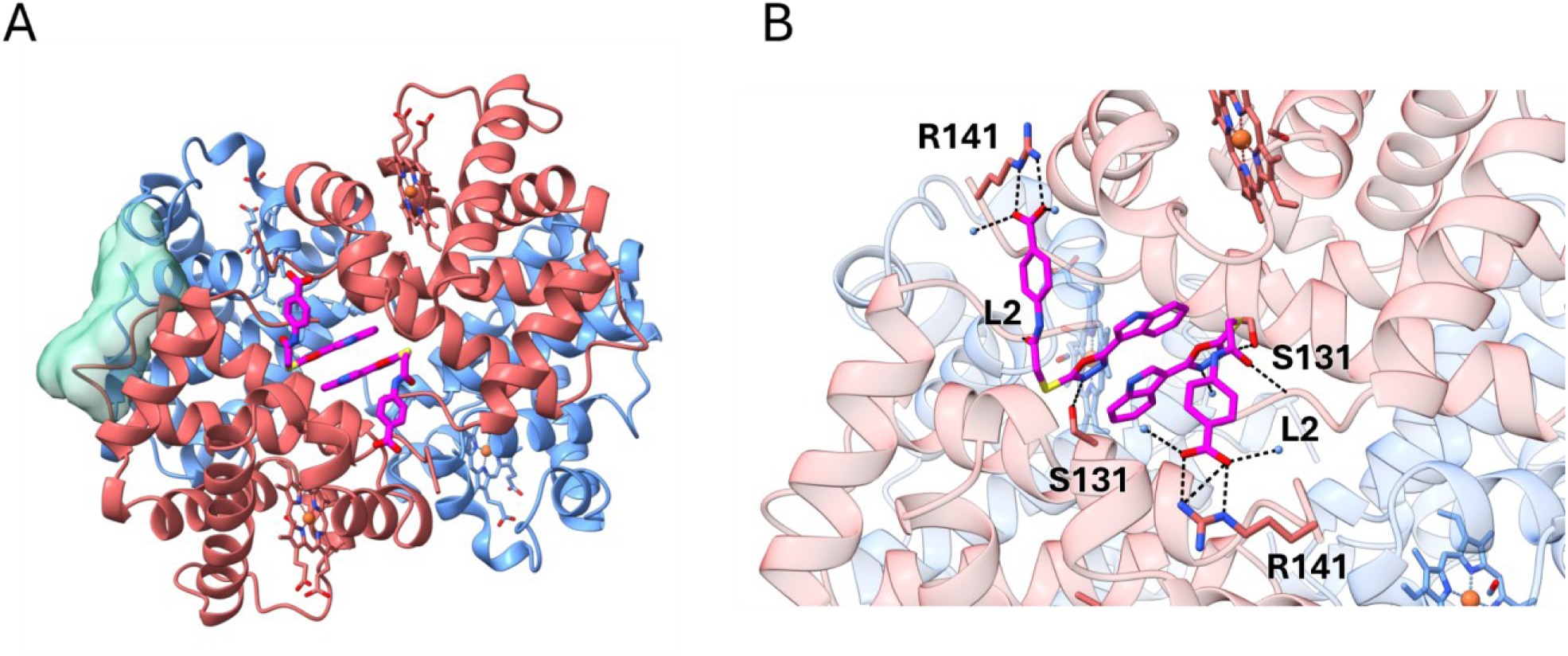
Binding of C35 to Hb. (**A)** The X-ray three-dimensional structure of Hb co-crystallized with the compound C35. Alpha subunits and beta subunits are shown as red and blue ribbons, respectively; the green surface represents the putative binding site of C35 as predicted by molecular docking [12]; C35 is shown in magenta. **(B)** Close-up of the binding site of C35 in the cleft between the two Hb alpha subunits (in red). Beta subunits are in blue, C35 in magenta, while contacts are shown as black dashed lines. Water molecules are represented as light blue spheres.

The binding stoichiometry of C35 to Hb observed in the crystals is 2:1, i.e. two molecules of C35 symmetrically bind one Hb tetramer, confirming the stoichiometry estimated by ITC [12]. However, the crystallographic structure identified a binding pocket that differs from the one proposed by docking in our recent paper [12] (Fig 2A). Indeed, the obtained structure shows C35 bound in a cleft between the Hb alpha subunits. In this pocket, the two C35 molecules form several symmetrical interactions with Hb alpha lining residues (Fig 2B). Specifically, the benzoic acid moiety of each C35 molecule interacts through a salt bridge with the positively charged lateral chain of Arg141. Each molecule of C35 establishes an H-bond between the oxadiazolic moiety and the side chain of Ser131 (helix H) and another H-bond between the amide group of C35 and the -NH on the backbone of Leu2. The oxadiazolic and the indolic moieties of both C35 molecules interact with each other through a stacking interaction. Furthermore, being the C35 binding site partially exposed to the solvent, different polar contacts with water molecules are also observed.

### C35 binds Hb in the same cleft of Hb left shifters and stabilizes a R2 conformation

Structural analysis of the Hb-C35 complex uncovered a previously unreported binding site for C35. To further investigate this new finding, we conducted a detailed comparison of Hb in complex with C35 with the structures of Hb deposited in the PDB.

This analysis was performed using the ‘’Structure similarity search’’ tool available on the PDB website (https://www.rcsb.org/search/advanced/structure). The tool performs a comprehensive analysis of the PDB, taking into account the entire protein assembly, including its ternary/quaternary state and any bound molecules. The first ten structures returned by the algorithm are listed in S2 Table and are all human Hb structures in the R conformation. The R state corresponds to a conformation of Hb with high oxygen affinity, where the α1β1 dimer is rotated by 15° with respect to the α2β2 dimer compared to the low oxygen affinity T state. Among the top 5 highest-scoring structures, two presented a ligand bound in a similar position with respect to C35 (i.e., PDB ID: 1QXE and PDB ID: 3IC0). In 1QXE Hb binds a molecule of 5-hydroxymethyl furfural (5-HMF) and in 3IC0 a derivative of vanillin (INN298). Both molecules were characterized by Safo’s group as Hb left-shifters [16,17], i.e., ligands that stabilize the R state with respect to the T state, thus increasing the oxygen affinity. These molecules have been developed as potential therapeutic agents against sickle cell anemia.

This finding raised the question of whether C35 might share similar properties and mode of action with those compounds. We focused on the comparison between C35 and INN298 (4-hydroxy-3-methoxybenzaldehyde), a more potent Hb left-shifter than 5-HMF (Fig 3A and Fig 3B).

**Fig 3.**
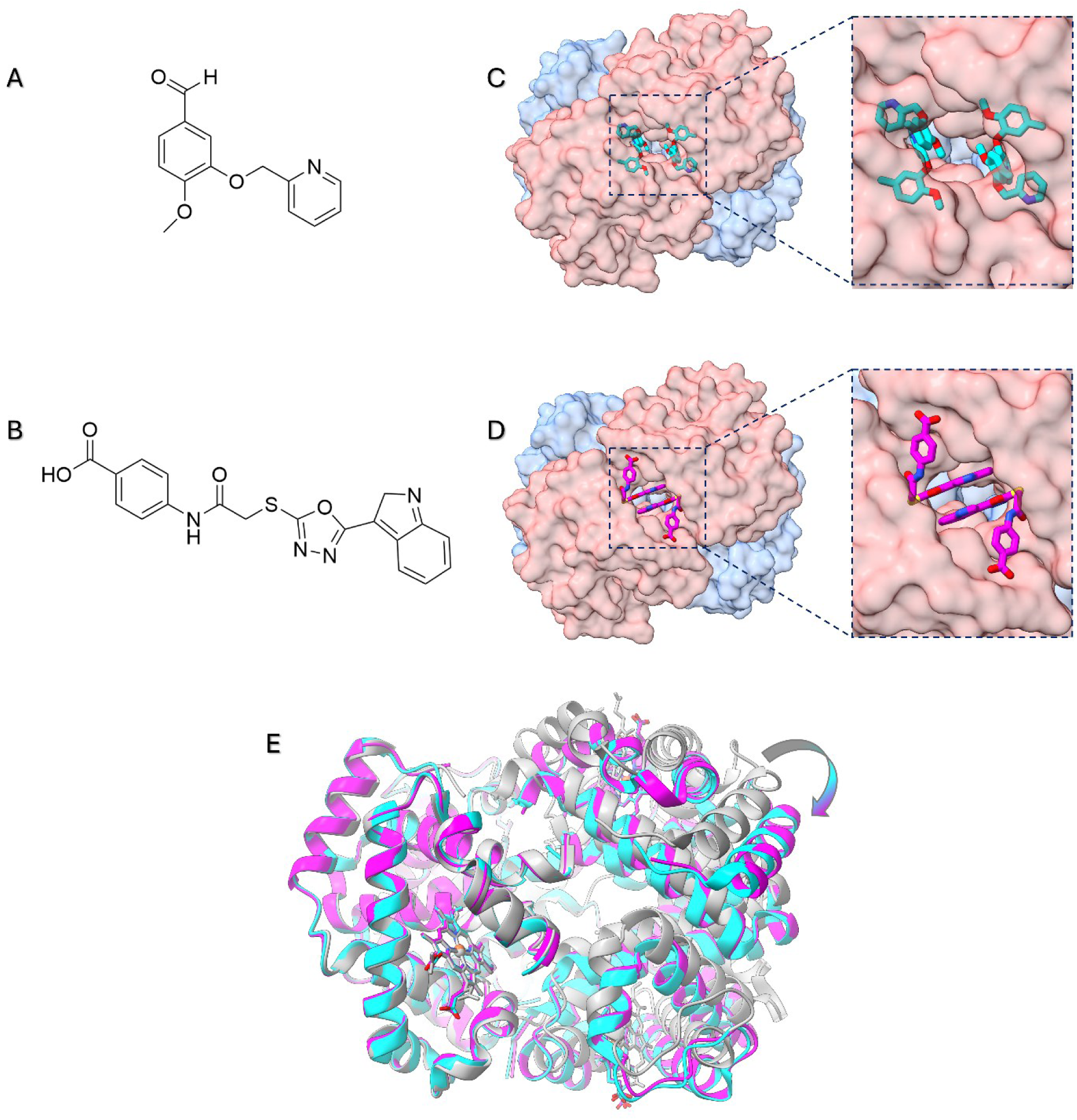
Structural comparison between Hb bound to C35 and Hb bound to INN298. (A) Chemical structure of the compound INN298. (B) Chemical structure of the compound C35. (C) Molecular surface of Hb bound to INN298 and its close-up on the INN298 binding site. (D) Molecular surface of Hb bound to C35 and its close-up on C35 binding site (PDB ID: 9HBA). INN298 and C35 are reported as cyan and magenta sticks, respectively. (E) Superimposition of the alpha-1 and beta-1 chains of different tetrameric Hb structures: HbCO (R state, PDB ID: 2DN3 – grey ribbons), Hb bound to INN298 (R2 state, PDB ID: 3IC0 – cyan ribbons) and Hb bound to C35 (R2 state, PDB ID: 9HBA – magenta ribbons). Heme molecules are represented as sticks. The RMSD values estimated for Hb bound to INN298 or to C35 considering R-state Hb as a reference are 4.46 Å and 4.39 Å, respectively. The RMSD value between INN298- and C35-bound Hbs is 0.54 Å.

INN298 and C35 share the same binding site on Hb (Figs 3C and 3D), although some differences are present. First, the binding stoichiometry of C35 is 2:1 (on tetramer base, *vide supra*), while the stoichiometry for INN298 is 4:1, with two of the four molecules forming a covalent bond with the terminal amino group of Val1 in the alpha chains, while the other two interact through non-covalent bonds. Differently from INN298, C35 only binds Hb *via* non-covalent interactions.

It is well known that compounds able to interact with Hb in the allosteric binding site between the alpha subunits can stabilize a relaxed R conformation. In particular, INN298 stabilizes an alternative quaternary Hb conformation named R2 state, over the R state trajectory [18,19]. The R and R2 states of tetrameric Hb mainly differ in the positioning of the α2β2 dimer relative to the α1β1 dimer. Therefore, to investigate the effect of C35 on the Hb quaternary structure, we compared C35-bound Hb with reference structures by calculating the global root mean square deviation (RMSD) on the α2β2 subunits after superimposing the α1β1 subunits. As references, we used R-state HbCO (PDB ID: 2DN3, [20]) and R2-state Hb bound to INN298 (PDB ID: 3IC0) (Fig 3E). The RMSD calculated relative to the R2 structure stabilized by INN298 is very low (0.54 Å) compared to that obtained relative to the R structure (4.39 Å). This result suggests that C35, similarly to INN298, can stabilize an R2 Hb conformation.

### Compound C35 increases Hb oxygen affinity

Considering that C35 and the left-shifter INN298 share the same binding site and induce comparable changes in Hb structure, we decided to compare the effect of the two molecules on the functional properties of Hb, i.e., on its oxygen binding affinity and cooperativity.

INN298, stabilizing the R conformation, reduces the binding cooperativity of oxygen to Hb and increases the oxygen affinity, thus also slowing down the HbS polymerization process that involves the deoxy form of the protein [13,17]. However, in the reference study [17], the effect of INN298 on oxygen binding was only investigated in intact RBCs, where, in addition to its direct action on Hb, the permeation of the effector into the cells must also be considered. Here, we decided to investigate the activity of INN298 at the highest concentration tested in [17] (i.e., 2 mM).

The reference oxygen binding curve of Hb showed a *p*50 of 8.1 ± 0.3 torr, with a Hill coefficient (n) of 1.9 ± 0.1 (Fig 4 and Table 1). Stripped Hb (i.e., Hb outside RBCs in the absence of any allosteric effector) has a reported n of approximately 2.4 [21]. The lower value measured here is likely due to the effect of DMSO (used at a concentration of 2% (v/v) to align with studies conducted with INN298 and C35), as reported in the work of Liu and colleagues [22]. As expected, INN298 shifted to the left the oxygen binding curve of Hb, with a *p*50 value of 0.8 ± 0.1 torr in the presence of the compound (Fig 4 and Table 1). Also, cooperativity was found to be strongly affected by the presence of INN298, with a Hill coefficient dropping to 0.8 ± 0.1.

**Fig 4.**
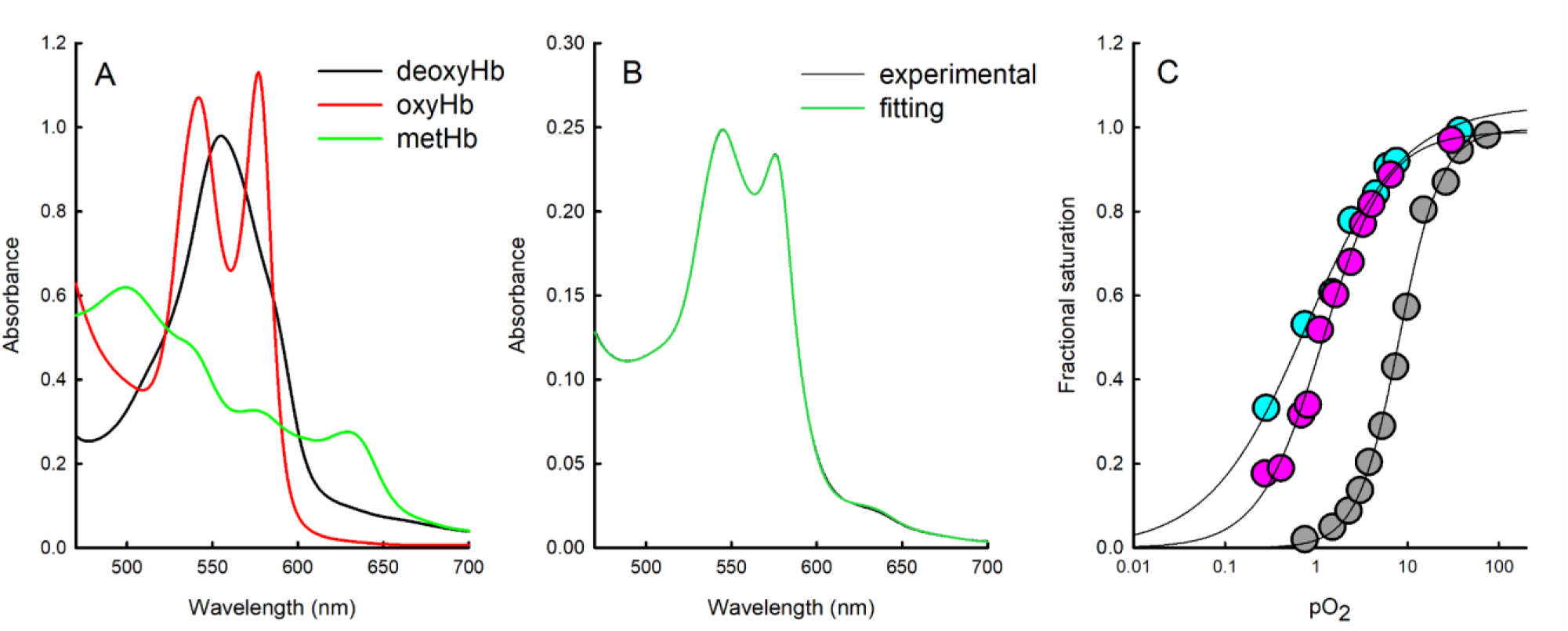
Effect of C35 and INN298 on the oxygen binding to Hb. (A) Reference spectra used for linear combination. (B) Example of comparison between an experimental spectrum (black line) measured at a pO2 of 9.4 torr and the corresponding fit to the linear combination of reference spectra (green line). (C) Oxygen binding curves of Hb measured at 37 °C in the absence (grey circles) and presence of either 1 mM C35 (magenta circles) or 2 mM INN298 (cyan circles). The data points represent the Hb fractional saturation with oxygen obtained from linear combination of reference spectra. Lines through data points are the fitting to the Hill equation (Eq. 3). The estimated values are reported in Table 1.

**Table 1.**
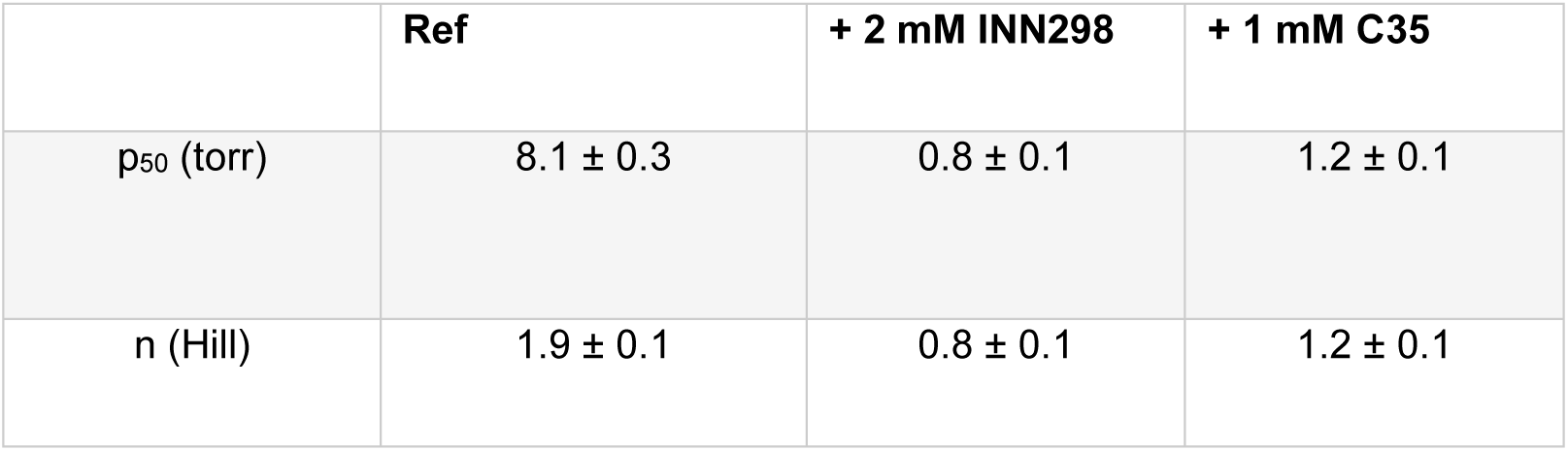
*p*50 and Hill coefficient of Hb in the absence and presence of either 2 mM INN298 or 1 mM C35 at pH 7.4, 37 °C.

|  | Ref | + 2 mM INN298 | + 1 mM C35 |
| --- | --- | --- | --- |
| $p_{50}$ (torr) | $8.1 \pm 0.3$ | $0.8 \pm 0.1$ | $1.2 \pm 0.1$ |
| n (Hill) | $1.9 \pm 0.1$ | $0.8 \pm 0.1$ | $1.2 \pm 0.1$ |

The oxygen binding curve in the presence of C35 was measured at a final concentration of 1 mM compound. This concentration is significantly higher than its dissociation constant (K_D_ = 0.57 ± 0.06 µM, [12]) but was necessary to saturate Hb, which was present at a concentration of 100 µM in the analysis to allow monitoring of the absorption of Q bands for the calculation of the fractional saturation (Fig 4A and 4B). The p50 calculated in the presence of C35 was 1.2 ± 0.1 torr (Fig 4C and Table 1), almost comparable to the value obtained in the presence of INN298. Also in this case, the cooperativity of oxygen binding is almost completely lost, with a Hill coefficient of about 1.2 ± 0.1 (Fig 4C and Table 1).

### C35 does not impair haptoglobin binding to Hb

In the crystal structure of tetrameric Hb with C35, the compound binds at the interface between the two Hb dimers, specifically at the level of the alpha subunits. Under physiological conditions, haptoglobin (Hp) binds free Hb dimers in the bloodstream *via* specific non-covalent interactions, forming a stable complex that prevents renal loss and oxidative damage by enabling Hb clearance through CD163-expressing macrophages [23]. During the initial virtual screening campaign, C35 was selected to interact with a region of Hb that would not interfere with Hp binding [12]. However, in its newly identified binding pose revealed by the crystal structure, the position of C35 is apparently not compatible with the co-presence of Hp bound to a Hb dimer. In fact, the structural alignment of a single Hb dimer bound to two C35 molecules (from the crystal structure of tetrameric Hb with C35) with the crystal structure of the Hb:Hp complex shows that C35 occupies a region involved in Hb:Hp interaction interface (Fig 5A). This observation prompted us to experimentally investigate whether C35 might affect Hb:Hp complex formation, as this potential interference would represent a serious limitation for the compound’s further development as a pharmacological candidate, either as antibiotic or Hb left-shifter.

**Fig 5.**
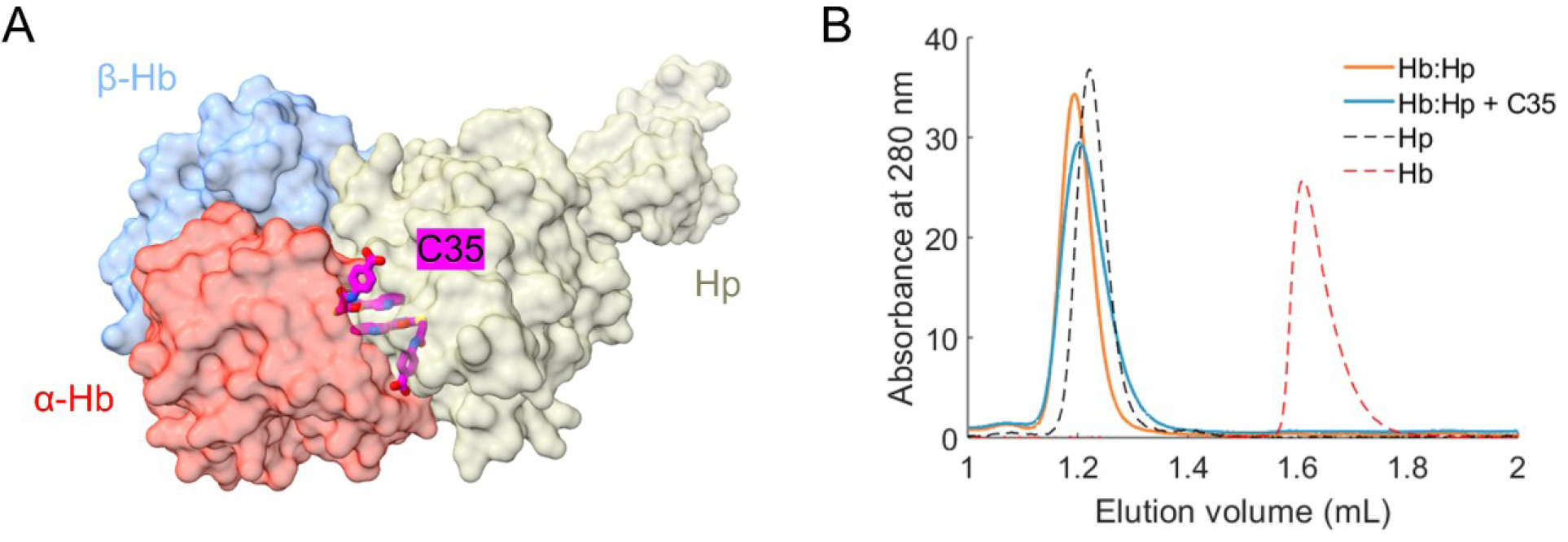
Effect of C35 binding on the formation of the Hb:Hp complex. (A) Comparison of the binding mode of C35 and Hp on a Hb dimer. A single Hb dimer of the crystal structure obtained in this study (PDB ID: 9HBA) was aligned with the structure of the Hb:Hp complex (PDB ID: 4F4O). Protein surfaces are shown as semi-transparent: Hb α-chains in red, Hb β-chains in blue, and a single Hp protomer (α- and β-chains) in yellow. C35 is shown as magenta sticks. (B) SEC profiles of isolated Hb (red dashed line) and Hp (black dashed line), and of the Hb:Hp complex (2:1 molar ratio; one Hb dimer per Hp protomer) in the absence (orange line) and presence (blue line) of 1 mM C35.

To explore the effect of C35 on Hb:Hp interaction, we performed SEC experiments. As a first step, experimental conditions were optimized to isolate the Hb:Hp complex chromatographically. Hb and Hp were mixed in solution at a 2:1 stoichiometric ratio (one Hb dimer per Hp protomer), and the resulting chromatogram was compared to those of the individual proteins run separately (Fig 5B). The chromatogram of the Hb-Hp mixture showed a single peak with a lower elution volume (1.19 mL) than each of the separate proteins (1.22 mL for Hp and 1.61 mL for Hb), indicating the formation of a higher molecular weight complex. Moreover, no residual peaks corresponding to the individual proteins were observed for the 2:1 Hb:Hp sample, confirming that the complex had been successfully isolated.

Maintaining the same protein concentrations and stoichiometry, we repeated the SEC experiment in the presence of C35, added at a concentration of 1 mM to both the sample and the mobile phase. While the resulting chromatographic peak appeared slightly broader compared to the control, the elution volume remained unchanged, and no additional peaks corresponding to unbound Hb or Hp were detected. These observations indicate that C35 does not disrupt the Hb:Hp complex under the tested conditions. Accordingly, no off-target effects related to Hb:Hp dissociation are expected for this compound.

## Discussion

In this work, we report the serendipitous identification of the small molecule C35 as a dual-function compound: an allosteric effector of Hb that increases its oxygen affinity, and a potent antimicrobial agent that inhibits *S. aureus* growth by interfering with iron acquisition. Originally selected for its ability to disrupt the Hb:IsdB interaction *in vitro*, C35 was shown to effectively inhibit *S. aureus* growth under iron-limited conditions, an effect that was reversed by excess of FeCl_3_, strongly suggesting iron starvation as the underlying antibacterial mechanism.

Structural studies revealed that C35 binds in a solvent-exposed cleft between the α-subunits of the Hb tetramer, a site shared with known Hb left-shifters such as INN298 [17] and Voxelotor [24]. This binding mode, distinct from the originally predicted docking pose, stabilizes an R2 quaternary conformation of Hb, characterized by high oxygen affinity and reduced cooperativity. Unlike most known left-shifters, which form covalent adducts with Hb, C35 interacts *via* non-covalent forces, representing a prototype molecule for a novel class of Hb modulators. Notably, this binding mode is, to date, shared only by PF-07059013, a non-covalent Hb left-shifter that recently entered clinical phase 1 trials and exhibits a similar interaction interface with nanomolar affinity for Hb (https://clinicaltrials.gov/study/NCT04323124). This convergence further underscores the growing pharmacological interest in non-covalent Hb modulators as safer and potentially more tunable therapeutic alternatives.

The inhibition of *S. aureus* growth in the presence of C35 raised the question of whether this activity could be linked to altered Hb oligomerization dynamics. Specifically, since heme extraction by the Isd system requires Hb dimerization [25], it was hypothesized that C35 may exert its inhibitory effect by stabilizing the tetrameric form of Hb, thereby limiting the accessibility of heme to IsdB. However, experimental analysis by SEC demonstrated that C35 does not significantly affect the tetramer-to-dimer equilibrium of Hb (S2 Fig), suggesting that the primary mechanism of action is not related to oligomeric stabilization but rather to direct competition with hemophore binding *via* conformational effects or steric hindrance.

It should be noted that Hb is expected to be mostly in the dimeric state at the concentrations used for the ELISA that was employed for the initial screening [12] and the *S. aureus* growth inhibition assay presented here. This hints to the possibility of C35 binding to the Hb dimer with a different (or a further) binding pose with respect to that observed for the tetramer in crystallographic experiments. The fact that Hb concentrations required for crystallization shift the dimer-tetramer equilibrium towards the latter hampers further investigation by crystallographic techniques.

To our knowledge, this is the first demonstration that a single ligand can simultaneously modulate oxygen binding to Hb, through the stabilization of a specific Hb quaternary structure, and interfere with heme scavenging by bacterial hemophores. This discovery opens new avenues for the development of dual-acting compounds that target both the functional modulation of Hb and bacterial iron acquisition. Rational design strategies could exploit C35’s scaffold to generate derivatives with controlled RBC permeability: cell-permeable analogs might serve as Hb left-shifters for the treatment of SCD, while impermeable variants could function extracellularly as antimicrobials during systemic infections. In terms of antimicrobial activity, moreover, it is worth noting that several bacterial iron acquisition systems rely on heme extraction from Hb [26,27]. Given that C35 exerts its effect through direct binding to Hb, it is conceivable that it could interfere with other bacterial iron acquisition systems that similarly exploit heme scavenging from host sources.

The clinical relevance of such dual-acting agents is underscored by the increased susceptibility of SCD patients to bacterial infections [28]. For this population, bifunctional C35 derivatives with balanced intra- and extracellular distribution could simultaneously address both pathological Hb polymerization and the associated infectious risk. Although the concentrations required to modulate intracellular and extracellular targets may differ substantially, it can be speculated that compounds favoring RBC uptake might still retain sufficient plasma levels to exert antibacterial activity, thereby offering a dual therapeutic benefit even with a predominant intracellular partitioning.

In conclusion, our study highlights the therapeutic potential of targeting the α-subunit interface of Hb as a novel pharmacological hotspot, offering a promising platform for the development of compounds that combine hemoglobinopathy treatment with antimicrobial action.

## Materials and Methods

### Chemicals and solubilization of C35

All reagents were purchased by MERCK (St. Louis, MO, USA) and were used as received. C35 was synthesized in-house, with a purity of at least 95%, as reported in [12] (see Scheme) and dissolved in 100% dimethyl sulfoxide (DMSO, Sigma-Aldrich) at a final concentration of 100 mM.

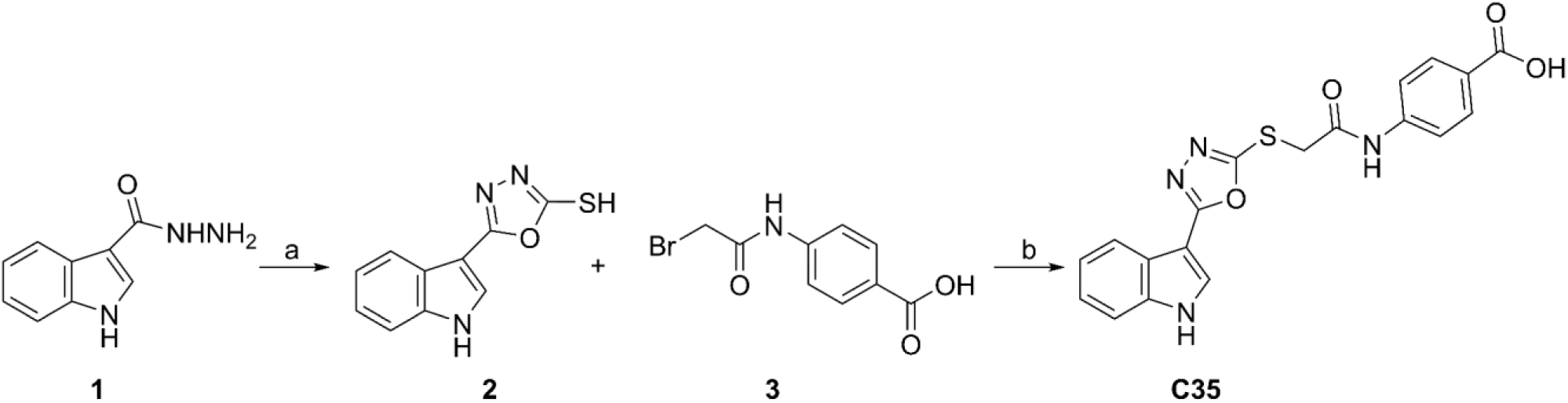

a. KOH, CS_2_, EtOH, reflux, 4 h. b) K_2_CO_3_, acetone reflux

### Bacterial strains and growth conditions

All experiments were carried out with *S. aureus* strain Newman [29] and its isogenic in frame-deletion mutant strain (Δ*isdB*). The Δ*isdB* mutant was obtained following the mutagenesis protocol described in Schuster et al. [30]. For inactivation of the *isdB* gene, a 955-bp fragment and a 989-bp fragment overlapping ATG and the TAA of *isdB*, respectively, were amplified by PCR using the primer couples UPFW*isdB*: 5’-AACTGCAGCCAAACCGTGTTAAACAATG-3’/UPRVi*sdB*: 5’-CCCAAGCTTGTTCATGTTGTAGAAACAAC-3’ and DWFW*isdB*: 5’-CCCAAGCTTAACTAATAAATCGTCTTTATATTT-3’/DWRV*isdB*:5’-CCGCTCGAGTGCTAGATTCACAAACGG-3’, respectively, and cloned in pIMAY*, yielding pIMAY*Δ*isdB*. Then, the latter was introduced by electroporation into *S. aureus* wild type. The functionality of the plasmid was then examined using a temperature-sensitivity test, confirming the ratio of colonies on plates incubated at 28 °C and 37 °C. Chromosomal integration *via* a single crossover event was achieved by incubating *S. aureus/*pIMAY*Δ*isdB* transformants at 37 °C with antibiotic selection and verifying by a colony PCR using the T3/T7 primers. The second crossing-over event was encouraged by culturing/sub-culturing the selected clones at 28 °C for at least 50 generations, followed by verification of plasmid loss using para-chlorophenylalanine (PCPA), a toxic phenylalanine analogue. Once chloramphenicol sensitivity was confirmed, genomic DNA was extracted from potential candidates and the deletion was confirmed by PCR using primers OUTFW*isdB*: 5’-TGTATACATAGGCGCAGACA-3’)/OUTRV*isdB*: 5’-AACTCGCGGTCTATTGCCA-3’, and PCR fragments were checked by sequencing.

Bacterial strains were routinely cultured in Tryptic Soy Agar (TSA) at 37 °C for 18 h and stored with 15% glycerol (v/v) at -80 °C. In this study, two media were used: *i)* Roswell Park Memorial Institute (RPMI-1640, Sigma-Aldrich) broth prepared according to the manufacturer’s instructions with a modification; RPMI was supplemented with 1% casamino acids (CAA) (w/v) to support bacterial growth, and *ii)* metal-depleted RPMI (NRPMI) prepared following the protocol described by Pishchany et al. [14]. Briefly, the RPMI medium was treated for 16 h at 4 °C with 70 g/L of the metal-chelating Chelex® resin (Bio-Rad) under moderate stirring, then filtered through Whatman no. 1 filter paper. The medium was subsequently supplemented with non-iron metals: 25 µM ZnCl_2_, 25 µM MnCl_2_, 100 µM CaCl_2_ and 1 mM MgCl_2_ prepared in advance as sterile 1,000x solutions. The pH of the NRPMI was then adjusted to 7.4, then sterilized by filtration using Corning® filter system (Corning) and stored at 4 °C.

When required, ethylenediamine-N,N’-bis(2-hydroxyphenylacetic acid) (EDDHA) was resuspended in anhydrous ethanol to a final concentration of 100 mM for sterilization, and then added to the culture medium at the required concentration. FeCl₃ (Sigma-Aldrich) was prepared as a 0.1 M stock solution in 10 mM HCl and stored at -20 °C. A stock solution of 930 µM human Hb (see below) was prepared and stored at -80 °C in phosphate-buffered saline (PBS). For C35, a 100 mM stock solution was freshly prepared in DMSO and stored at -20 °C.

The ability of *S. aureus* WT and its isogenic Δ*isdB* to grow in NRPMI was investigated by monitoring bacterial growth (OD_600_) over time, as described in Pishchany et al. [14]. For that, *S. aureus* WT and its isogenic Δ*isdB* mutant were pre-cultured in RPMI supplemented with 500 µM EDDHA, to restrict iron availability, for 16-20 h at 37 °C with 180 rpm shaking. The next day, cultures were centrifuged for 5 min at 7,500 *x g* and the pellets were resuspended in NRPMI containing 500 µM EDDHA and normalized to OD_600_ = 3. 10 µL of bacterial suspension were subcultured into 1 mL NRPMI in a 15 ml screw-cap conical tube, supplemented with either 500 µM EDDHA and/or 80 nM Hb and/or 100 µM C35, and incubated at 37 °C with vigorous shaking. Bacterial growth (OD_600_) was measured in a multiplate reader (SPARK 10M TECAN®) every 6-12 h for up to 96 h.

### Protein purification

Human hemoglobin A (Hb) was purified from outdated blood donated by non-smoking volunteers to a blood transfusion center, following the procedure described in Viappiani et al. [31]. Written informed consent was obtained from all donors, and both the blood donation and the use of outdated samples complied with Italian law 219/2005 on blood donation and usage. RBCs were washed with saline and lysed under hypotonic conditions by adding seven volumes of Buffer Hb1 (10 mM HEPES pH 6.9, 1 mM EDTA). The lysate was clarified by centrifugation (23,000 *x g*, 1 h, 4 °C), and the supernatant containing oxygenated Hb was dialyzed against Buffer Hb1. The sample was then loaded onto a CM-Sephadex C-50 column (100 × 5 cm). Soluble RBC components were separated from Hb using a linear gradient from 0% to 80% Buffer Hb2 (10 mM HEPES pH 8.6, 1 mM EDTA); Hb was subsequently eluted using a gradient from 80% to 85% Buffer Hb2. The purified Hb was dialyzed into storage buffer (10 mM HEPES pH 7.2, 1 mM EDTA), aliquoted, flash-frozen in liquid nitrogen, and stored at −80 °C. The concentration and oxidation state of oxyHb were assessed by UV–Vis absorption spectroscopy using known molar extinction coefficients for heme-specific absorbance peaks [32].

Strep-tagged IsdB was expressed and purified as described previously [33]. Strep-tag® II-IsdB (residues 125–485, UniProt Q8NX66) was codon-optimized for *Escherichia coli*, cloned into pASK-IBA3-plus vector, and expressed in *E. coli* BL21 strain induced with 0.2 μg/mL anhydrotetracycline at 20 °C for 20 h. Cells were lysed, and the protein purified using Strep-Tactin®XT affinity resin followed by SEC in Buffer W. Final yield exceeded 100 mg/L with >95% purity. Protein concentration was calculated using ε_280nm_ = 47,790 M⁻¹ cm⁻¹, with holo-IsdB content estimated to be <5% based on heme absorbance at 405 nm (ε_405nm_ = 90,500 M⁻¹ cm⁻¹) [33].

### Hb co-crystallization with C35

Crystals of carboxyhemoglobin (HbCO) grew in the presence of 10 mM C35 in a solution containing 3.0 M ammonium sulfate and 1% (w/v) 2-methyl-2,4-pentanediol (MPD) in sitting drop conditions after which the crystals were cryo-cooled in liquid nitrogen for data collection. X-ray diffraction data was collected at Diamond Light Source beamline i03 at wavelength 0.9763Å. Data was integrated and scaled using the CCP4 package; structures were solved by molecular replacement using Phaser crystallographic software [34]. The structural model was iteratively refined and rebuilt by using the Coot program [35]. Ligand coordinates and restraints were generated from the SMILES string using the Grade software package (Global Phasing Ltd). The structure coordinates are deposited on Protein Data Bank (PDB) (9HBA), and the data collection and refinement statistics are shown in S1 Table.

To investigate the presence of global 3D-shape similarity, using the C35-bound Hb as a reference, the ‘’Structure similarity search’’ tool on the PDB website (https://www.rcsb.org/search/advanced/structure) was exploited. The search was conducted by uploading the C35-bound Hb structure in pdb format without relaxation step and considering only molecular assemblies.

### Oxygen binding to Hb

Oxygen-binding curves of 100 µM Hb were measured with a modified tonometer [36] in 100 mM HEPES, 1 mM EDTA, pH 7.4 at 37 ± 0.4 °C in either the presence or absence of studied effectors. The oxygenated Hb (oxyHb) concentration was estimated using the extinction coefficient at 415 nm of 125,000 M^−1^ cm^−1^. The Hayashi reducing system was exploited to prevent autoxidation [37] during measurements. Before measurement, oxyHb was deoxygenated under helium (50 mL/min) for 60 minutes to obtain the deoxygenated form (deoxyHb). Samples were then exposed to different partial oxygen pressures (pO_2_), generated using an Environics 4000 gas mixer (Environics Inc, Tolland, CT, U.S.A.), connected to a helium bottle and different premixed helium/oxygen bottles. Spectra were collected after 30 minutes incubation time in the 450–700 nm range using a Cary 4000 spectrophotometer (Agilent, Santa Clara, CA, USA). The control experiments, performed in the absence of any ligands, were carried out in the presence of 2% DMSO (v/v).

The Hb fractional saturation at each pO_2_ was calculated by adapting the method reported by Rivetti et al. [38]. Spectra were analyzed as a linear combination of the deoxyHb, oxyHb and metHb spectra (i.e., the reference spectra) measured under the same experimental conditions (Eq. 1):

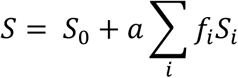

Where *S* is the measured absorbance spectrum, *S_0_* is an offset, *a* is the scale factor to correct for the intensity of the analyzed spectrum, the index *i* denotes the heme species (oxyheme, deoxyheme and metheme), *S_i_* are the reference spectra and *f_i_* are the fractional coefficients. The estimation of the oxygen saturation (y) is obtained as (Eq. 2):

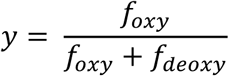

The pO_2_ corresponding to 50% fractional saturation (*p*50) and the Hill coefficient (n) were estimated by fitting the calculated fractional saturation at different pO_2_s to the Hill equation (Eq. 3):

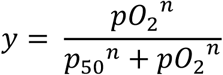

Where *y* is the oxygen saturation, *a* is the amplitude, *x* is the pO_2_, *p*50 is the oxygen partial pressure corresponding to 0.5 fractional saturation and *n* is the Hill coefficient, accounting for binding cooperativity.

### Isothermal Titration Calorimetry

Experiments were performed at 25 °C using 12 µM IsdB and 1 mM C35 in a buffered solution containing 50 mM HEPES buffer, pH 7.6. C35 was first dissolved at 100 mM in DMSO and then diluted in the final buffer to a concentration of 1 mM. Solutions were degassed for 10 min under vacuum before the titration. DMSO was added to the IsdB solution at a final concentration of 1% to balance the amount of DMSO in the C35 solution. ITC titrations were carried out using a MicroCal PEAQ-ITC instrument (Malvern, Malvern, UK). C35 was added to the instrument measurement cell, containing 280 μL of IsdB, by a first addition of 0.4 μL and 18 subsequent additions of 2 μL. A time interval of 150 s was set between the addition of each aliquot of C35. To subtract the dilution heat, a reference experiment was performed in which the reaction cell was filled only with the buffer solution with 1% DMSO, while the syringe was filled with 1 mM C35. Experiments were performed under continuous stirring at 750 rpm. All experiments were performed in triplicate. Data analysis was performed using MicroCal PEAQ-ITC Analysis Software (version 1.41, Malvern Panalytical, Malvern, UK).

### Size-exclusion chromatography

SEC was performed to assess whether compound C35 exerts a destabilizing effect on the Hb:haptoglobin (Hb:Hp) complex and/or on the quaternary structure of isolated Hb (i.e., its tetrameric form).

The destabilizing effect of compound C35 on the Hb:Hp complex was investigated by exploiting an Agilent 1260 HPLC system (Agilent Technologies, Inc., Santa Clara, CA, USA) equipped with a Superdex® 200 Increase 3.2/300 column (Cytiva). The column was pre-equilibrated with PBS buffer (pH 7.4) either in the absence or presence of 1 mM C35, and operated at a flow rate of 0.07 mL/min. Samples containing Hb, Hp (Phenotype 1-1) or the Hb:Hp complex (prepared using a 2:1 stoichiometry, i.e., one Hb dimer associated with one Hp protomer), at concentrations of 2 mg/mL, were injected and eluted at 25 °C. Elution profiles were monitored by measuring absorbance at 280 nm and 406 nm using an Agilent 1260 Infinity II WR Diode Array Detector.

The estimation of the dissociation constant for the Hb dimer/tetramer equilibrium was carried out using an ÄKTA Pure 25 M chromatographic system (GE Health Sciences™, Chicago, IL, USA) equipped with a Superdex 75 Increase 5/150 GL column (GE Health Sciences™) with a mobile phase consisting of 20 mM Tris, 150 mM NaCl and 1 mM EDTA pH 8.0 at 20 °C, either in the absence or presence of 0.01 mM C35. Control experiments in the absence of C35 were carried out adding 0.01% (v/v) DMSO. The amount of DMSO in the control experiments accounts for the same amount of DMSO related to C35-treated samples. The flow rate was set to 0.3 mL/min. The separation was run at room temperature and the absorbance of the column effluent was monitored both at 280 nm (for calibration) and to 415 nm (for Hb samples). Hb was loaded at the following concentrations: 0.2, 1, 5, 15 and 30 µM. The calibration curve was built using conalbumin (75 kDa), ovalbumin (43 kDa) and lysozyme (14 kDa) as standard proteins (Cytiva). The concentrations reported in S2 Fig also consider column dilution, which was approximately 3-fold with respect to the loading concentrations. Data points in S2 Fig were fitted using (Eq. 4):

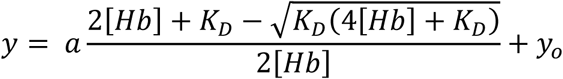

where *y* is the estimated apparent molecular weight, *a* is the amplitude, corresponding to the apparent molecular weight of the tetramer, [Hb] is the Hb concentration, *K_D_* is the dissociation constant for the Hb dimer/tetramer equilibrium and *y_0_* is an offset, corresponding to the apparent molecular weight of the Hb dimer.

## Acknowledgements

We thank Maisem Laabei, University of Bristol, UK, for kindly providing *Staphylococcus aureus* Newman and the pIMAY* plasmid. We thank Gianmarco Mangiaterra (University of Urbino Carlo Bo, Urbino, Italy) for technical assistance in constructing the pIMAY*Δ*isdB* plasmid. We thank Dr. Davide Cavazzini (Department of Chemistry, Life Sciences and Environmental Sustainability, University of Parma) for technical support in size-exclusion chromatography experiments. This work is dedicated to the memory of Prof. Andrea Mozzarelli, full professor of Biochemistry at the University of Parma who sadly passed away on August 17^th^, 2024. We are most grateful to Prof. Mozzarelli for his seminal work on hemoglobin allosteric regulation and kinetic mechanisms of sickling that inspired many of us and a plethora of students throughout his scientific and academic career.

## Author Contribution

SB, EF, LR, ODB, MM, FS and FM designed the project and conceptualized the approach; SB, EF, BC, LR, FS, LL and SF supervised the project; SB, FS and EF provided financial resources; EG, SA, BR, FS and LL designed, synthesized and purified C35 compound for assays; SH performed the microbiological assays and analyzed data; FM, MM, VBC and ODB performed the biochemical characterization and analyzed data; CC, VBC and SF performed the ITC measurements and analyzed data; PB performed protein crystallization and structural analysis; SH, FM, MM, VBC, EG, SA, CC and ODB prepared the figures; FM, SB, BC and ODB wrote the original draft; all authors have read and agreed to the published version of the manuscript.

## Data availability

The dataset generated and analyzed in this study has been deposited in the Figshare database under accession code https://doi.org/10.6084/m9.figshare.29145080. Crystallographic model presented in this study is available under accession code 9HBA.

## Supporting information

**S1 Fig. Characterization of the binding of C35 to IsdB.** Raw data for ITC titration of 12 µM IsdB with 1 mM C35 (top panel); binding isotherm of the integrated titration curve (bottom panel). The experiment was carried out at 25 °C in 50 mM HEPES buffer, pH 7.6.

**S2 Fig. Effect of C35 on the tetramer/dimer equilibrium of oxygenated Hb.** (A) Dependence of the apparent molecular weight as a function of Hb concentrations both in the absence (white circles) and presence of 0.01 mM C35 (magenta circles). Data points in the absence of C35 are obtained in the presence of the same DMSO concentration (i.e., 0.1% v/v) with respect to data points obtained in the presence of C35. The fitting of the data (solid lines) to Eq. 3 allowed for the estimation of dissociation constants of about 0.67 ± 0.15 and 0.80 ± 0.38 µM for the Hb in the absence and presence of C35, respectively, indicating that the compound does not significantly alter the tetramer-to-dimer equilibrium of Hb. (B) Chromatograms of calibrants (lower panel) used to build the calibration curve (upper panel); CONA, OVA and LISO correspond to conalbumin, ovalbumin and lysozyme, respectively.

**S1 Table.** Crystal data and structure refinements for Hb:C35 complex.

**S2 Table.** PDB ID, score and corresponding DOI of the first ten structures obtained exploiting “Structure similarity search’’ tool on the PDB website (https://www.rcsb.org/search/advanced/structure) and using the structure of Hb bound to C35 as a reference.

## Notes

### Competing Interest Statement

The authors have declared no competing interest.

### Summary of Updates

This version of the manuscript has not been revised. However, we have added the supplementary materials and included the ORCID identifiers for all authors.

### Funding

- Ministry of Education, Universities and Research: 2020AE3LTA
- Ministry of Education, Universities and Research
- Ministry of Education, Universities and Research: PNRR MUR Â-M4C2Â–Investimento 1.4Â–Avviso Â“Centri NazionaliÂ”Â–D.D. n. 3138 del 16 dicembre 2021
- University of Turin: SPYF_RILO_01_23, LAZL_RILO_01_23

## References

1. Sati H, Carrara E, Savoldi A, Hansen P, Garlasco J, Campagnaro E, et al. The WHO Bacterial Priority Pathogens List 2024: a prioritisation study to guide research, development, and public health strategies against antimicrobial resistance. Lancet Infect Dis. 2025. doi:10.1016/S1473-3099(25)00118-5

2. Marchetti M, De Bei O, Bettati S, Campanini B, Kovachka S, Gianquinto E, et al. Iron metabolism at the interface between host and pathogen: From nutritional immunity to antibacterial development. Int J Mol Sci. 2020;21(6):2145. doi:10.3390/ijms21062145

3. Mazmanian SK, Skaar EP, Gaspar AH, Humayun M, Gornicki P, Jelenska J, et al. Passage of Heme-Iron Across the Envelope of Staphylococcus aureus. Science. 2003;299:906–9. doi:10.1126/science.1081147

4. De Bei O, Marchetti M, Ronda L, Gianquinto E, Lazzarato L, Chirgadze DY, et al. Cryo-EM structures of staphylococcal IsdB bound to human hemoglobin reveal the process of heme extraction. Proc Natl Acad Sci U S A. 2022;119:e2116708119. doi:10.1073/pnas.2116708119

5. Andrade MA, Ciccarelli FD, Perez-Iratxeta C, Bork P. NEAT: a domain duplicated in genes near the components of a putative Fe3+siderophore transporter from Gram-positive pathogenic bacteria. Genome Biol. 2002;3:research0047.1. doi:10.1186/gb-2002-3-9-research0047

6. Torres VJ, Pishchany G, Humayun M, Schneewind O, Skaar EP. Staphylococcus aureus IsdB is a hemoglobin receptor required for heme iron utilization. J Bacteriol. 2006;188:8421–9. doi:10.1128/JB.01335-06

7. Pietrocola G, Pellegrini A, Alfeo MJ, Marchese L, Foster TJ, Speziale P. The iron-regulated surface determinant B (IsdB) protein from Staphylococcus aureus acts as a receptor for the host protein vitronectin. J Biol Chem. 2020;295:10008–22. doi:10.1074/jbc.RA120.013510

8. Waryah CB, Gogoi-Tiwari J, Wells K, Eto KY, Masoumi E, Costantino P, et al. Diversity of virulence factors associated with west australian methicillin-sensitive staphylococcus aureus isolates of human origin. Biomed Res Int. 2016;2016:8651918. doi:10.1155/2016/8651918

9. Gonzalez JJI, Hossain MF, Neef J, Zwack EE, Tsai C-M, Raafat D, et al. TLR4 sensing of IsdB of Staphylococcus aureus induces a proinflammatory cytokine response via the NLRP3-caspase-1 inflammasome cascade. mBio. 2024;15. doi:10.1128/mbio.00225-23

10. Tsai C-M, Caldera JR, Hajam IA, Chiang AWT, Tsai C-H, Li H, et al. Non-protective immune imprint underlies failure of Staphylococcus aureus IsdB vaccine. Cell Host Microbe. 2022;30:1163–72.e6. doi:10.1016/j.chom.2022.06.006

11. Skaar EP, Schneewind O. Iron-regulated surface determinants (Isd) of Staphylococcus aureus: Stealing iron from heme. Microbes Infect. 2004;6:390–7. doi:10.1016/j.micinf.2003.12.008

12. Cozzi M, Failla M, Gianquinto E, Kovachka S, Buoli Comani V, Compari C, et al. Identification of small molecules affecting the interaction between human hemoglobin and Staphylococcus aureus IsdB hemophore. Sci Rep. 2024;14:8272. doi:10.1038/s41598-024-55931-8

13. Mozzarelli A, Hofrichter J, Eaton WA. Delay Time of Hemoglobin S Polymerization Prevents Most Cells from Sickling in Vivo. Science. 1987;237:500–6. doi:10.1126/science.3603036

14. Pishchany G, Haley KP, Skaar EP. Staphylococcus aureus Growth using Human Hemoglobin as an Iron Source. J Vis Exp. 2013. doi:10.3791/50072

15. Hammer ND, Skaar EP. Powerful genetic resource for the study of methicillin-resistant Staphylococcus aureus. mBio. 2013. doi:10.1128/mBio.00166-13

16. Abdulmalik O, Safo MK, Chen Q, Yang J, Brugnara C, Ohene-Frempong K, et al. 5-Hydroxymethyl-2-furfural modifies intracellular sickle haemoglobin and inhibits sickling of red blood cells. Br J Haematol. 2005;128:851–8. doi:10.1111/j.1365-2141.2004.05332.x

17. Abdulmalik O, Ghatge MS, Musayev FN, Parikh A, Chen Q, Yang J, et al. Erratum: Crystallographic analysis of human hemoglobin elucidates the structural basis of the potent and dual antisickling activity of pyridyl derivatives of vanillin (Acta Crystallographica Section D (2011) D67 (920-928)). Acta Crystallogr D Biol Crystallogr. 2011;67:929. doi:10.1107/S0907444911045860

18. Silva MM, Rogers PH, Arnone A. A third quaternary structure of human hemoglobin A at 1.7-Å resolution. J Biol Chem. 1992;267:17248–56. doi:10.1016/s0021-9258(18)41919-9

19. Schumacher MA, Zheleznova EE, Poundstone KS, Kluger R, Jones RT, Brennan RG. Allosteric intermediates indicate R2 is the liganded hemoglobin end state. Proc Natl Acad Sci U S A. 1997;94:7841–6. doi:10.1073/pnas.94.15.7841

20. Park SY, Yokoyama T, Shibayama N, Shiro Y, Tame JRH. 1.25 Å Resolution Crystal Structures of Human Haemoglobin in the Oxy, Deoxy and Carbonmonoxy Forms. J Mol Biol. 2006;360:683–96. doi:10.1016/j.jmb.2006.05.036

21. Caccia D, Ronda L, Frassi R, Perrella M, Del Favero E, Bruno S, et al. PEGylation promotes hemoglobin tetramer dissociation. Bioconjug Chem. 2009;20:1546–55. doi:10.1021/bc900130f

22. Liu C, Bo A, Cheng G, Lin X, Dong S. Characterization of the structural and functional changes of hemoglobin in dimethyl sulfoxide by spectroscopic techniques. Biochim Biophys Acta. 1998;1385:181–9. doi:10.1016/S0167-4838(98)00044-2

23. Andersen CBF, Stødkilde K, Sæderup KL, Kuhlee A, Raunser S, Graversen JH, et al. Haptoglobin. Antioxid Redox Signal. 2017;26:814–31. doi:10.1089/ars.2016.6793

24. Glaros AK, Razvi R, Shah N, Zaidi AU. Voxelotor: alteration of sickle cell disease pathophysiology by a first-in-class polymerization inhibitor. Ther Adv Hematol. 2021;12:20406207211001136. doi:10.1177/20406207211001136

25. De Bei O, Marchetti M, Guglielmo S, Gianquinto E, Spyrakis F, Campanini B, et al. Time-resolved X-ray solution scattering unveils the events leading to hemoglobin heme capture by staphylococcal IsdB. Nat Commun. 2025;16:1361. doi:10.1038/s41467-024-54949-w

26. Kronstad JW, Caza M. Shared and distinct mechanisms of iron acquisition by bacterial and fungal pathogens of humans. Front Cell Infect Microbiol. 2013;3:80. doi:10.3389/fcimb.2013.00080

27. Akinbosede D, Chizea R, Hare SA. Pirates of the haemoglobin. Microb Cell. 2022;9:60–75. doi:10.15698/MIC2022.04.775

28. Abdulmanea AA, Alharbi NS, Somily AM, Khaled JM, Algahtani FH. The Prevalence of the Virulence Genes of Staphylococcus aureus in Sickle Cell Disease Patients at KSUMC, Riyadh, Saudi Arabia. Antibiotics. 2023;12(7):1221. doi:10.3390/antibiotics12071221

29. Baba T, Bae T, Schneewind O, Takeuchi F, Hiramatsu K. Genome Sequence of Staphylococcus aureus Strain Newman and Comparative Analysis of Staphylococcal Genomes: Polymorphism and Evolution of Two Major Pathogenicity Islands. J Bacteriol. 2008;190:300–10. doi:10.1128/JB.01000-07

30. Schuster CF, Howard SA, Gründling A. Use of the counter selectable marker pheS* for genome engineering in staphylococcus aureus. Microbiology (Reading). 2019;165:1020–30. doi:10.1099/mic.0.000791

31. Viappiani C, Abbruzzetti S, Ronda L, Bettati S, Henry ER, Mozzarelli A, et al. Experimental basis for a new allosteric model for multisubunit proteins. Proc Natl Acad Sci U S A. 2014;111:12758–63. doi:10.1073/pnas.1413566111

32. Reichlin M. Enzyme Proteins: Hemoglobin and Myoglobin in Their Reactions with Ligands . Eraldo Antonini and Maurizio Brunori. North-Holland, Amsterdam, 1971 (U.S. distributor, Elsevier, New York). xx, 436 pp., illus. $30. Frontiers of Biology, vol. 21. . Science. 1972;178:296. doi:10.1126/science.178.4058.296

33. Gianquinto E, Moscetti I, De Bei O, Campanini B, Marchetti M, Luque FJ, et al. Interaction of human hemoglobin and semi-hemoglobins with the Staphylococcus aureus hemophore IsdB: a kinetic and mechanistic insight. Sci Rep. 2019;9:18944. doi:10.1038/s41598-019-54970-w

34. McCoy AJ, Grosse-Kunstleve RW, Adams PD, Winn MD, Storoni LC, Read RJ. Phaser crystallographic software. J Appl Crystallogr. 2007;40:658–74. doi:10.1107/S0021889807021206

35. Emsley P, Lohkamp B, Scott WG, Cowtan K. Features and development of Coot. Acta Crystallogr D Biol Crystallogr. 2010;66:486–501. doi:10.1107/S0907444910007493

36. Ronda L, Bruno S, Abbruzzetti S, Viappiani C, Bettati S. Ligand reactivity and allosteric regulation of hemoglobin-based oxygen carriers. Biochim Biophys Acta Proteins Proteomics. 2008;1784:1365–77. doi:10.1016/j.bbapap.2008.04.021

37. Hayashi A, Suzuki T, Shin M. An enzymic reduction system for metmyoglobin and methemoglobin, and its application to functional studies of oxygen carriers. Biochim Biophys Acta. 1973;310:306–16. doi:10.1016/0005-2795(73)90110-4

38. Rivetti C, Mozzarelli A, Rossi GL, Henry ER, Eaton WA. Oxygen Binding by Single Crystals of Hemoglobin. Biochemistry. 1993;32:2888–900. doi:10.1021/bi00062a021

