## Supporting information for "Two birds with one stone: a novel potential antibiotic blocking IsdB-mediated heme extraction by *Staphylococcus aureus* with serendipitous hemoglobin left-shifting activity"

¶ Contributed equally to the work

* Corresponding authors

**S1 Fig. Characterization of the binding of C35 to IsdB.** Raw data for ITC titration of 12 µM IsdB with 1 mM C35 (top panel); binding isotherm of the integrated titration curve (bottom panel). The experiment was carried out at 25 °C in 50 mM HEPES buffer, pH 7.6.

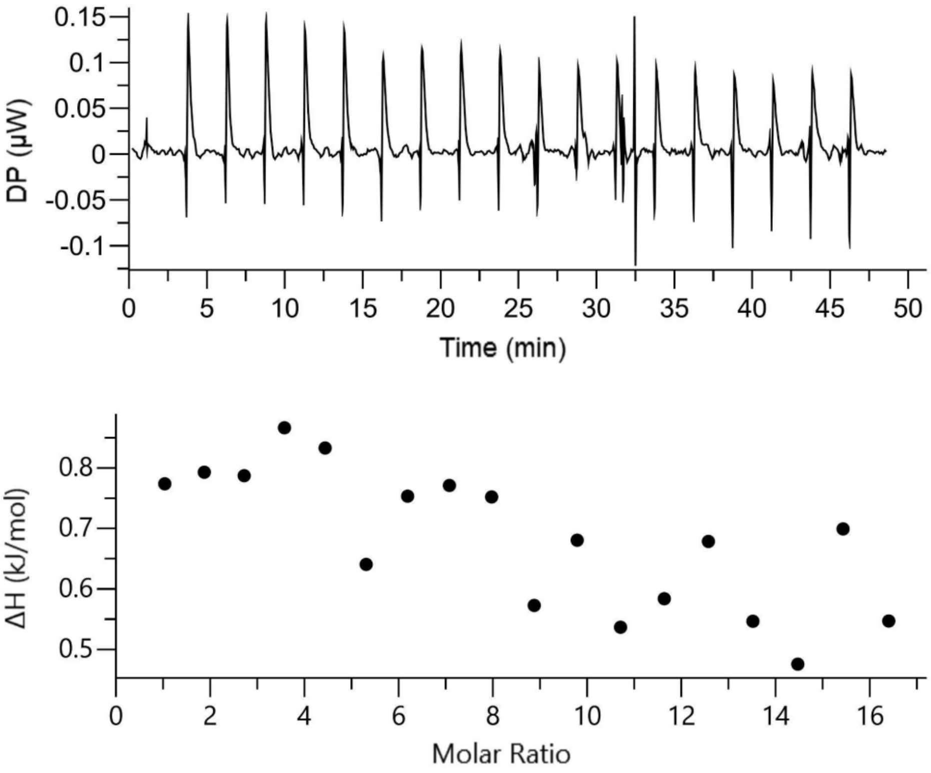

**S2 Fig. Effect of C35 on the tetramer/dimer equilibrium of oxygenated Hb.** (A) Dependence of the apparent molecular weight as a function of Hb concentrations both in the absence (white circles) and presence of 0.01 mM C35 (magenta circles). Data points in the absence of C35 are obtained in the presence of the same DMSO concentration (i.e., 0.1% v/v) with respect to data points obtained in the presence of C35. The fitting of the data (solid lines) to Eq. 3 allowed for the estimation of dissociation constants of about 0.67 ± 0.15 and 0.80 ± 0.38 μM for the Hb in the absence and presence of C35, respectively, indicating that the compound does not significantly alter the tetramer-to-dimer equilibrium of Hb. (B) Chromatograms of calibrants (lower panel) used to build the calibration curve (upper panel); CONA, OVA and LISO correspond to conalbumin, ovalbumin and lysozyme, respectively.

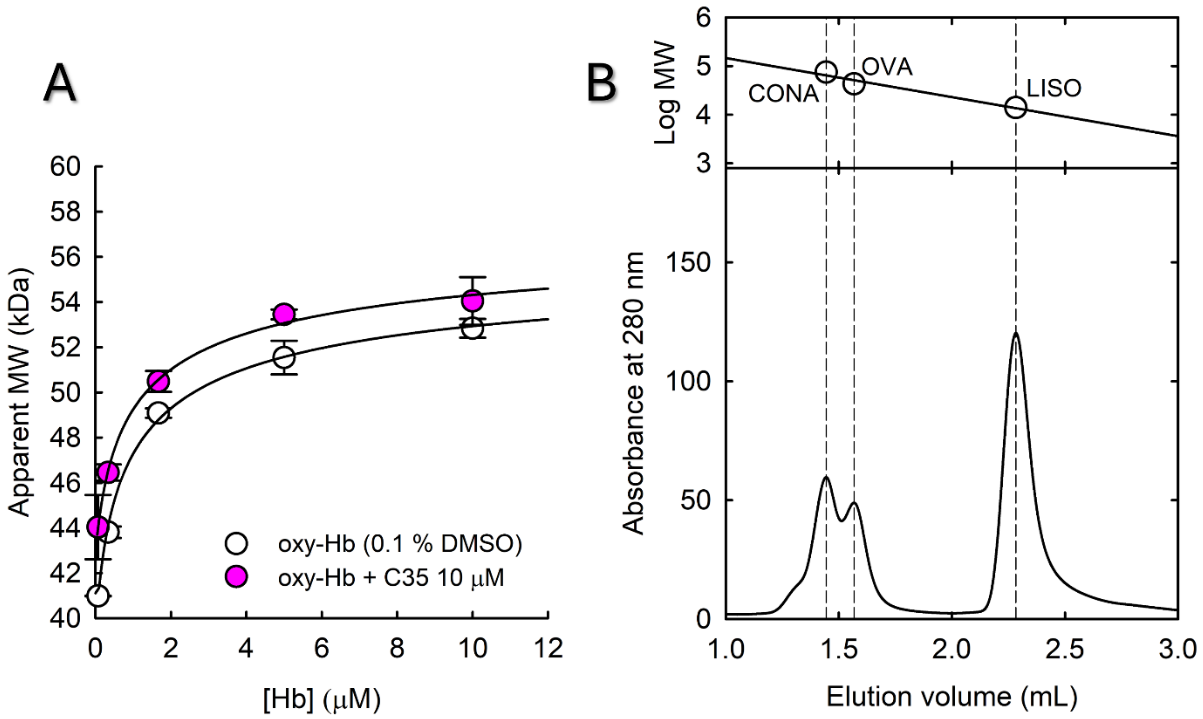

**S1 Table.** Crystal data and structure refinements for Hb:C35 complex.

| item | 9HBA |
|---|---|
| Collection date | 29/01/2023 |
| Data Collection: | |
| Beamline | DIAMOND BEAMLINE I03 |
| Wavelength | 0.9763 |
| Resolution range | 53.26 - 1.51 (1.540 - 1.510) |
| Space group | P 32 2 1 |
| Cell (a b c) | 92.18 92.18 142.97 |
| Cell (alpha beta gamma) | 90.00 90.00 120.00 |
| Total reflections | 110596 |
| Unique reflections | 110596 (5481) |
| Multiplicity | 20.1 (14.6) |
| Completeness (%) | 100.0 (99.4) |
| Mean I/sigma(I) | 14.5 (0.3) |
| R-merge | 0.095 (4.40) |
| R-pim | 0.022 (1.18) |
| CC-half | 1.000 (0.32) |
| Refinement: | |
| Refinement resolution range | 53.26 - 1.51 (1.55 - 1.51) |
| No. reflections | 109875 (8051) |
| No. reflections (Rfree) | 5451 (368) |
| R-factor | 0.184 (0.437) |
| Rfree | 0.215 (0.472) |
| Number of total atoms | 5098 |
| atoms for macromolecules | 4363 |
| atoms for ligands | 412 |
| atoms for waters | 323 |
| Average B-factor | 31.2 |
| RMS(bonds) | 0.01 |
| RMS(bond angles) | 1.555 |
| RMS(dihedral angles) | 5.286 |
| Crystallisation conditions | 3.0 M Ammonium sulfate; 1% (w/v) MPD |
| Ligand SMILES | O=C(CSc1nnc(-c2c[nH]c3ccccc23)o1)Nc1ccc(C(=O)[O-])cc1 |

**S2 Table.** PDB ID, score and corresponding DOI of the first ten structures obtained exploiting "Structure similarity search'' tool on the PDB website (https://www.rcsb.org/search/advanced/structure) and using the structure of Hb bound to C35 as a reference.

| PDB ID | Rank | Structure Match Score | Description | Ligand | Organism |
|---|---|---|---|---|---|
| 6KAS | 1 | 87.22 | Carbonmonoxy human hemoglobin A in the R2 quaternary structure at 95 K: Dark | No ligand | Homo sapiens |
| 1QXE | 2 | 87.20 | Structural Basis for the Potent Antisickling Effect of a Novel Class of 5-Membered Heterocyclic Aldehydic Compounds | 5-hydroxymethyl-furfural | Homo sapiens |
| 6L5X | 3 | 86.82 | Carbonmonoxy human hemoglobin A in the R2 quaternary structure at 95 K: Light (2 min) | No ligand | Homo sapiens |
| 3IC0 | 4 | 86.50 | Crystal Structure of liganded hemoglobin in complex with a potent antisickling agent, INN-298 | INN-298 | Homo sapiens |
| 6KAT | 5 | 86.18 | Carbonmonoxy human hemoglobin A in the R2 quaternary structure at 95 K: Light | No ligand | Homo sapiens |
| 6KAU | 6 | 86.07 | Carbonmonoxy human hemoglobin A in the R2 quaternary structure at 140 K: Dark | No ligand | Homo sapiens |
| 6L5Y | 7 | 85.06 | Carbonmonoxy human hemoglobin A in the R2 quaternary structure at 140 K: Light (2 min) | No ligand | Homo sapiens |
| 6KAV | 8 | 84.93 | Carbonmonoxy human hemoglobin A in the R2 quaternary structure at 140 K: Light | No ligand | Homo sapiens |
| 1QXD | 9 | 84.60 | Structural Basis for the Potent Antisickling Effect of a Novel Class of 5-Membered Heterocyclic Aldehydic Compounds | Furfural | Homo sapiens |
| 3IC2 | 10 | 84.27 | Crystal Structure of liganded hemoglobin in complex with a potent antisickling agent, INN-266 | INN-266 | Homo sapiens |
